# Pavian: Interactive analysis of metagenomics data for microbiomics and pathogen identification

**DOI:** 10.1101/084715

**Authors:** Florian P. Breitwieser, Steven L. Salzberg

## Abstract

**Summary:** Pavian is a web application for exploring metagenomics classification results, with a special focus on infectious disease diagnosis. Pinpointing pathogens in metagenomics classification results is often complicated by host and laboratory contaminants as well as many non-pathogenic microbiota. With Pavian, researchers can analyze, display and transform results from the Kraken and Centrifuge classifiers using interactive tables, heatmaps and flow diagrams. Pavian also provides an alignment viewer for validation of matches to a particular genome.

**Availability and implementation:** Pavian is implemented in the R language and based on the Shiny framework. It can be hosted on Windows, Mac OS X and Linux systems, and used with any contemporary web browser. It is freely available under a GPL-3 license from http://github.com/fbreitwieser/pavian. Furthermore a Docker image is provided at https://hub.docker.com/r/florianbw/pavian.

**Supplementary information:** Supplementary data is available at Bioinformatics online.

## Introduction

Metagenomics sequencing has the potential to revolutionize pathogen detection in infectious diseases. Currently, most infectious disease diagnosis is done with traditional culture-based methods, which are time consuming and labor intensive, and can miss off-target pathogens. Several recent studies showcased how DNA and RNA sequencing of patient samples can guide clinicians to identify the correct pathogenic species, even when standard methods are inconclusive (e.g. Wilson *et al.*, 2014; Salzberg *et al.*, 2015). The general workflow is to extract DNA or RNA from a patient sample that may be either fresh or preserved, to prepare a sequencing library, and then to generate millions or tens of millions of reads. Various bioinformatics tools are available to classify the reads; i.e., to label each read with the identify of a species (e.g. Lindgreen *et al.*, 2016). While the tools can demonstrate highly accurate classification results on simulated sequencing data, the interpretation of the real data is often difficult due to contamination both in the samples (Salter *et al.*, 2014) and in reference databases (Merchant *et al.*, 2014). Furthermore, negative controls or biological replicates are often unavailable for clinical samples. When looking for evidence of a pathogen in a sample that is a complex mixture of human DNA, nonpathogenic DNA, and a small amount of pathogenic DNA, interactive exploration and visualization of the data can help to find the needle in the haystack.

With Pavian, we provide a novel interface to explore metagenomics results from Kraken (Wood and Salzberg, 2014), Centrifuge (Kim *et al.*, 2016) and MetaPhlAn (Truong *et al.*, 2015) classifiers. While powerful tools for interactive microbiome analysis are available, such as Shiny-phyloseq (McMurdie and Holmes, 2015) and Seed (Beck *et al.*, 2015), they are geared towards ecological and community analysis. On the other hand, specialized pipelines for pathogen detection (e.g. Naccache *et al.*, 2014, Byrd *et al.* (2014)) do not provide an interactive way to explore the data. Compared to Taxonomer (Flygare *et al.*, 2016), which integrates read classification and visualization, Pavian provides additional ways of visualization and comparison of several samples. Pavian enables a researcher to dissect single samples as well to compare identifications across multiple samples. This interface aims to simplify the interpretation of the large, often complex results from metagenomics sequencing experiments.

## Pavian Software

Pavian implements a straightforward interface to analyze and compare complex metagenomics datasets. It can be installed easily on workstations or deployed on remote servers. The data shown in Figure 1 and used as the basis of the walk-through of the interface in supplementary file 1 is from a recent study in which we used metagenomic sequencing to identify pathogens in brain biopsies of patients with suspected neurological infections (Salzberg *et al.*, 2016).

**Figure 1:**
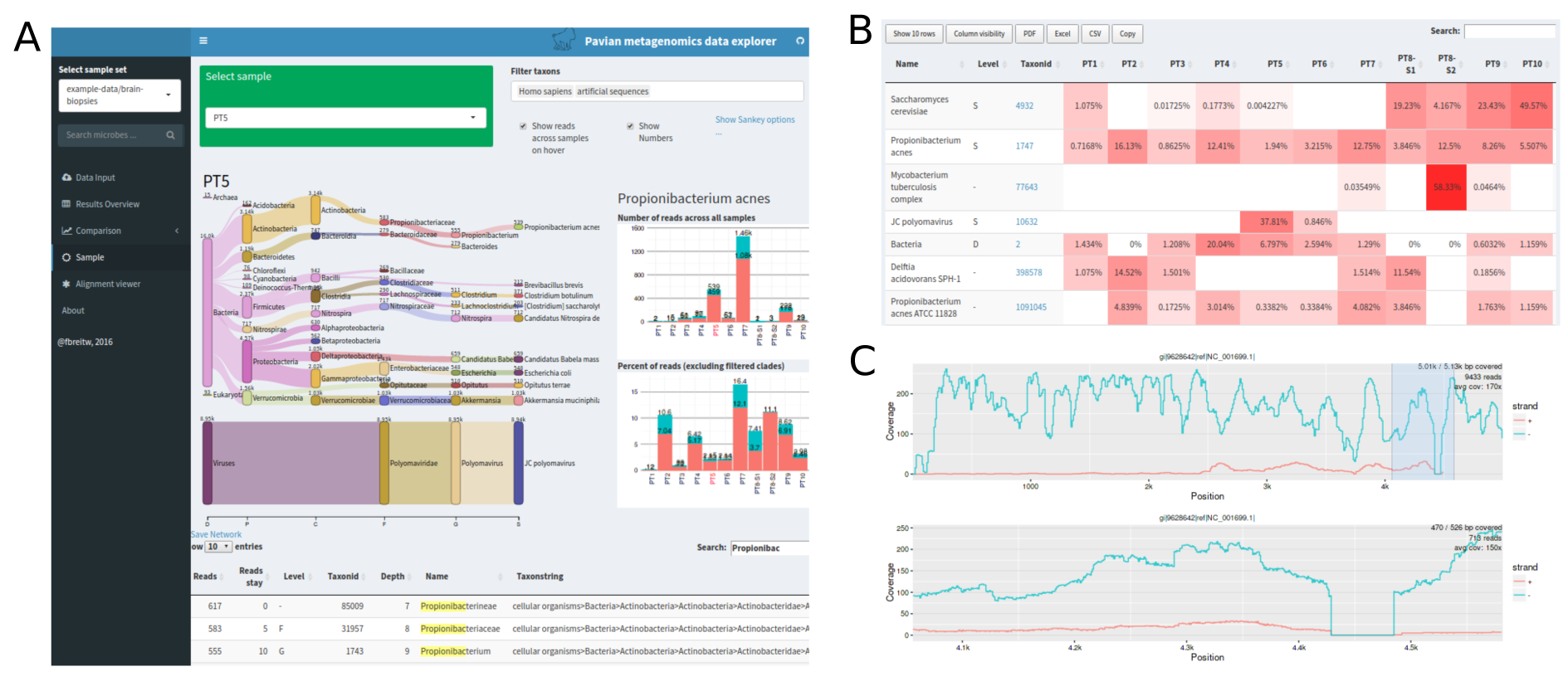
(A) Main interface of Pavian showing the side bar options and the identifications in sample PT5 in a Sankey diagram. The width of the flow corresponds to the number of reads, and on hover a barchart with the distribution of reads in other samples is presented. (B) Sample comparison interface. (C) Interactive alignment viewer.

**Sample view**: The sample view provides an overview of the classifications (see Figure 1A). Pavian's default visualization choice is a Sankey diagram, a type of graph that is often used to map energy flows (e.g. Subramanyam *et al.*, 2015). Pavian uses this diagram to display the flow of reads from the root of the taxonomy to more specific levels. The width of the flow is proportional to the number of reads. The Sankey diagram puts a visual emphasis on the major flows within the system and thus provides a clear visual summary of the classifications. Less interesting clades may be filtered out interactively.

**Sample comparison**: The sample comparison view juxtaposes the identification results from multiple samples (Figure 1B). This allows users to identify which microbes are commonly observed and which are present only in one or a few samples. The main view is a query-able table with taxa as rows and samples as columns. The third column provides an inline barchart representation of the data across all samples. By default, read counts at all taxonomy levels are shown, but it is possible to show only specific taxonomical levels. The data can be transformed by total sum scaling, variance-stabilizing transformation, and a robust z-score statistic. The results can also be displayed as an interactive heatmap.

**Alignment viewer**: A high read count for a particular species does not always mean that the microbe is present– contaminated genomes and low-complexity sequences can produce spurious assignments. By re-aligning the reads to the genome in question we can assure that the reads are uniformly distributed across the chromosomes. The built-in viewer takes bam format files from standard read aligner such as bowtie2 or bwa-mem (Figure 1C). The interface further provides links to download RefSeq genome assemblies.

**Implementation and installation**: Pavian is written in R using the Shiny framework (R Core Team, 2016, Chang *et al.* (2016)). It incorporates several interactive plots in Javascript and D3 (Bostock *et al.*, 2011). The tables are created using the R library DT / Javascript library DataTables, and use AJAX to transmit data on demand. The Sankey diagram is based on D3's sankey code (Bostock *et al.*, 2011) and the R library networkD3. The alignment viewer uses the library Rsamtools to load the bam file. While the backend can be launched from an R environment or installed on a server, the interface is accessed through a web browser. Thus the only requirement for viewing samples is a recent version of a web browser in any operating system. No expert knowledge is needed to use the interface.

To install and run Pavian within R, type

~~~
devtools::install_github('fbreitwieser/pavian') ## requires library devtools
pavian::**runApp**(port=5000)
~~~

For even easier installation, we also provide a Docker container in the public repository ‘florianbw/pavian’ at DockerHub. On Windows and Mac OSX, Docker Toolbox provides a convenient interface (https://www.docker.com/products/docker-toolbox). When using the command line version of docker, type

~~~
docker pull 'florianbw/pavian'
docker run–rm–p 5000:80 florianbw/pavian
~~~

In both cases, the interface will be available at http://127.0.0.1:5000 in your browser.

## Discussion

For assembled metagenomics data and binning, we highly recommend Anvi'o (Eren *et al.*, 2015).

## Conclusion

Pavian is a novel tool for visualizing and analyzing metagenomics data. Its functionalities helps microbiome researchers as well as clinical microbiologists to gain a better understanding of their data. It is freely available at https://github.com/fbreitwieser/pavian.

## Acknowledgments

We thank Daehwan Kim, Thomas Mehoke, Peter Thielen, and David Karig for helpful conversations and discussions.

*Funding*: This research was supported in part by NIH under grants R01 HG006677 and R01 GM083873, and by the U. S. Army Research Office under grant W911NF-14-1-0490.

